# Machine learning identifies large-scale reward-related activity modulated by dopaminergic enhancement in major depression

**DOI:** 10.1101/669887

**Authors:** Yuelu Liu, Roee Admon, Emily L. Belleau, Roselinde H. Kaiser, Rachel Clegg, Miranda Beltzer, Franziska Goer, Gordana Vitaliano, Parvez Ahammad, Diego A. Pizzagalli

## Abstract

**Background:** Theoretical models have emphasized systems-level abnormalities in Major Depressive Disorder (MDD). For unbiased yet rigorous evaluations of pathophysiological mechanisms underlying MDD, it is critically important to develop data-driven approaches that harness whole-brain data to classify MDD and evaluate possible normalizing effects of targeted interventions. Here, using an experimental therapeutics approach coupled with machine-learning we investigated the effect of a pharmacological challenge aiming to enhance dopaminergic signaling on whole-brain’s response to reward-related stimuli in MDD.

**Methods:** Using a double-blind placebo-controlled design, functional magnetic resonance imaging (fMRI) data from 31 unmedicated MDD participants receiving a single dose of 50 mg amisulpride (MDD_Amisulpride_), 26 MDD participants receiving placebo (MDD_Placebo_), and 28 healthy controls receiving placebo (HC_Placebo_) were analyzed. An importance-guided machine learning technique for model selection was used on whole-brain fMRI data probing reward anticipation and consumption to identify features linked to MDD (MDD_Placebo_ vs. HC_Placebo_) and dopaminergic enhancement (MDD_Amisulpride_ vs. MDD_Placebo_).

**Results:** Highly predictive classification models emerged that distinguished MDD_Placebo_ from HC_Placebo_ (AUC=0.87) and MDD_Placebo_ from MDD_Amisulpride_ (AUC=0.89). Although reward-related striatal activation and connectivity were among the most predictive features, the best truncated models based on whole-brain features were significantly better relative to models trained using striatal features only.

**Conclusions:** Results indicate that, in MDD, enhanced dopaminergic signaling restores abnormal activation and connectivity in a widespread network of regions. These findings provide new insights into the pathophysiology of MDD and pharmacological mechanism of antidepressants at the system level in addressing reward processing deficits among depressed individuals.

**ClinicalTrials.gov identifier:** NCT01253421 and NCT01701258

## Introduction

Major depressive disorder (MDD) is a debilitating disorder, often characterized by anhedonia (1), which is poorly addressed by current treatments (1, 2). Converging evidence across species suggests that mesocorticolimbic dopaminergic pathways involving the striatum are essential for reward processing (3–5). Dysfunction in this circuit has been associated with deficits in reward processing across psychiatric diseases (6). In MDD, neuroimaging studies have documented decreased striatal activation and reduced functional connectivity between the striatum and other nodes of the brain reward system in response to reward-related stimuli (7–9). Notably, some of these abnormalities were found to be acutely restored by pharmacologically-induced dopaminergic enhancement (10).

Despite advancements in our understanding of the pathophysiology of MDD, an unresolved issue is how enhanced dopaminergic signaling might modulate large-scale whole-brain activation and functional coordination in MDD. Besides the striatum, other brain regions, including the orbitofrontal cortex, amygdala, and anterior cingulate cortex, have been implicated in reward processing (11–14). Given that antidepressant treatments aiming to increase dopaminergic signaling might have faster therapeutic onsets (15, 16), it is important to investigate the effects of dopaminergic enhancement to gain a more thorough understanding of the potential neural mechanism through which these interventions may address reward processing deficits in MDD. Equally important, for an unbiased yet rigorous evaluation of pathophysiological mechanisms underlying MDD, it is critically important to develop data-driven approaches that harness whole-brain data to classify individuals with vs. without MDD and evaluate putative normalization of MDD-related abnormalities.

Towards this goal, we used a machine learning based approach to analyze whole-brain functional magnetic resonance imaging (fMRI) data collected from a double-blind placebo-controlled study, in which unmedicated individuals with MDD and healthy controls (HCs) performed a monetary incentive delay (MID) task after being randomized to either a single low dose of amisulpride (50 mg) or placebo. Amisulpride, a selective dopamine D2/D3 receptor antagonist, was selected because of its high affinity to block presynaptic autoreceptors at lower doses, thereby increasing dopamine release (17). In a first step, to identify the effects of enhanced dopaminergic transmission on reward-related brain activity, whole-brain fMRI data were entered into an importance-guided model selection procedure (based on the logistic regression with elastic net regularization; Fig. 1) to identify brain regions in which reward-related metrics were most predictive of differences between the MDD individuals receiving amisulpride vs. placebo. Next, to investigate the potential normalizing effect of enhanced dopaminergic transmission on MDD-related abnormalities, brain regions from the above step were compared with those most predictive of differences between MDD and HC group receiving placebo. To further assess putative normalization, in secondary analyses, we compared MDD individuals randomized to amisulpride vs. HC receiving placebo. Based on prior findings (7,10,18–22), we hypothesized that (1) under placebo, MDD would be associated with widespread reward-related abnormalities along the brain’s reward pathway and (2) transient DA enhancement would rescue such abnormalities.

**Figure 1:**
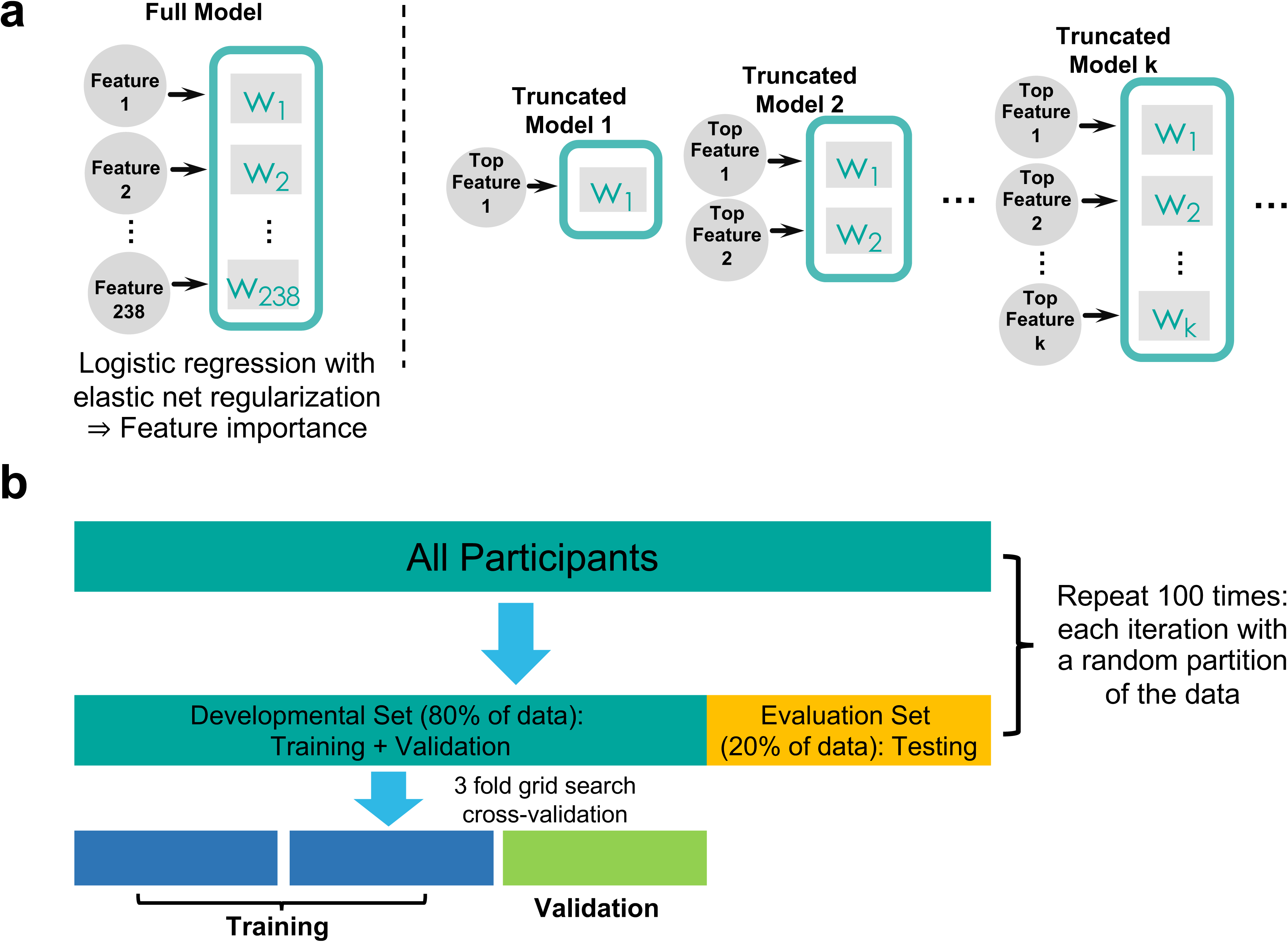
**a)** An illustration of the importance-guided sequential model selection procedure used to find the optimal set of features. First, a full model including all features is trained using logistic regression with elastic net regularization to determine relative importance of individual features. Next, a series of truncated models were trained based on a progressively increasing set of top features rank ordered by the full model. The set of features in the best parsimonious model on the evaluation set were deemed as the optimal feature set. **b)** An illustration of the nested cross-validation procedure used to train, validate, and test the models. A grid search procedure with 3-fold cross-validation was implemented on the developmental set to determine the best model parameters. The resulting model was further tested on the evaluation set, which contained an independent set of participants *not* used in training and validation. The entire procedure was repeated on 100 different random partitioning of the data to allow for stable model performance.

## Methods and Materials

### Participants

Participants were recruited by the Center for Depression, Anxiety and Stress Research at McLean Hospital using online advertisements, mailing and flyers within the Boston metropolitan areas for two independent studies using identical procedures.

Across the first (ClinicalTrials.gov identifier: NCT01253421) and second (NCT01701258) study, 62 unmedicated individuals with MDD (34 randomized to amisulpride, 28 randomized to placebo) and 63 demographically-matched healthy controls (placebo: N=30, amisulpride: N=33) were run in the imaging session. For the current analyses, we focused on analyses aiming at classifying case vs. controls (MDD_Placebo_ vs. HC_Placebo_ model) and classify the potential normalizing effects of dopaminergic enhancement (MDD_Placebo_ vs. MDD_Amisulpride_ model); thus, 92 participants were considered. Among these 92, 85 had useable fMRI data. Note that a subset of these participants (46 MDD, 23 randomized to amisulpride, 23 to placebo; 20 HC controls randomized to placebo) were included in a recent study that used a region-of-interest (ROI) approach to probe the effects of MDD and amisulpride on striatal activation and functional connectivity (10). Groups were matched for age, gender, ethnicity, and years of education (Table 1). General inclusion criteria were: right-handedness, age between 18–45, no MRI contraindications, no lifetime substance dependence, no past-year substance abuse, and no serious medical conditions. For the MDD groups, a diagnosis of MDD according to the Structured Clinical Interview for DSM-IV-TR Axis I Disorders (SCID) (2) was required, and exclusion criteria included: psychotropic medication in the past 2 weeks (6 weeks for fluoxetine, 6 months for dopaminergic drugs or antipsychotics) and any other axis I disorders (however, social anxiety disorder, simple phobia, or generalized anxiety disorder were allowed if secondary to MDD). For HC, exclusion criteria were: any medication in the last 3 weeks, current or past psychiatric illnesses (SCID), and first-degree familial psychiatric illness. Participants received $15/hour in addition to earnings in the fMRI task. The two protocols were approved by Partners Human Research Committee, and all participants provided written informed consent.

**Table 1:** Clinical and demographic characteristics of the participants.

| Characteristic | MDD<sub>Amisulpride</sub> |  | MDD<sub>Placebo</sub> |  | HC<sub>Placebo</sub> |  |
| --- | --- | --- | --- | --- | --- | --- |
|  | N = 31 |  | N = 26 |  | N = 28 |  |
|  | Mean | SD | Mean | SD | Mean | SD |
| Age (years) | 27.2 | 7.7 | 25.6 | 5.0 | 25.1 | 6.1 |
| Education (years) | 15.4 | 2.2 | 16.8 | 3.0 | 15.2 | 2.9 |
| Beck Depression Inventory 2<sup>nd</sup> Ed. | 26.3 | 7.9 | 26.7 | 7.9 | 1.8 | 2.7 |
| Hamilton Depression Rating Scale | 15.6 | 3.7 | 16.7 | 5.3 | 1.0 | 1.2 |
| Mood and Anxiety Symptom Questionnaire |  |  |  |  |  |  |
| Total Score | 168.5 | 22.9 | 174.1 | 21.7 | 91.5 | 13.3 |
| General Distress Anxiety Subscore | 23.6 | 5.1 | 25.4 | 6.6 | 12.3 | 1.2 |
| General Distress Depression Subscore | 37.9 | 9.4 | 39.0 | 9.3 | 13.9 | 2.0 |
| Anxious Arousal Subscore | 24.0 | 6.0 | 25.6 | 6.4 | 18.4 | 2.0 |
| Anhedonic Depression Subscore | 82.9 | 11.2 | 84.1 | 9.1 | 47.0 | 11.3 |
| Snaith-Hamilton Pleasure Scale | 31.7 | 4.7 | 31.4 | 7.0 | 22.8 | 6.7 |
| Duration of Current Major Depressive Episode (months) | 17.3 | 20.0 | 17.6 | 31.9 | N/A | N/A |
| Number of Past Depressive Episodes | 3.2 | 2.6 | 3.3 | 3.2 | N/A | N/A |
|  | N | % | N | % | N | % |
| Female | 28 | 90.3 | 19 | 73.1 | 22 | 81.5 |
| Caucasian | 20 | 64.5 | 13 | 50.0 | 13 | 48.1 |
| Current Comorbid Anxiety Disorders | 10 | 32.3 | 11 | 42.3 | N/A | N/A |
| Past Comorbid Anxiety Disorders | 13 | 41.9 | 12 | 46.2 | N/A | N/A |

Note: Groups were matched for age, gender, race, and years of education (one-way ANOVA; $\chi^2$ -test). All participants were right-handed. Between the MDD<sub>Amisulpride</sub> and MDD<sub>Placebo</sub> group, participants were matched for current and past comorbid anxiety disorders, as well as clinical scale measures ( $\chi^2$ -test; two-sample t-test).

### Procedure

The two studies followed identical procedures, pharmacological challenge, and MRI acquisition. In the first session, a PhD- or Masters-level clinician administered the SCID to determine eligibility, and participants filled out self-report scales (Table 1 and *Supplement*). In the second session, participants performed the MID task during fMRI scanning after receiving a single dose of amisulpride or placebo. The MID task was started one hour after pill administration due to pharmacokinetic data indicating that plasma concentration of amisulpride has a first peak approximately 1–1.5 hours after administration (17).

### fMRI Task

The MID has been described in detail (10, 23). Briefly, the task includes anticipation and receipt of monetary rewards (and penalties), which robustly recruit mesocorticolimbic regions (12, 13) and has been used to uncover reward-related abnormalities in both magnitude of activation and functional connectivity in MDD (7, 9, 10, 22, 24).

### Data Acquisition and Preprocessing

For both studies, MRI data were acquired at the McLean Imaging Center using a Siemens Tim Trio 3T MR scanner equipped with a 32-channel head coil. See *Supplementary Methods* for acquisition parameters and preprocessing.

### Feature Extraction

The features used in our classifiers consisted of coefficients from the single-subject level general linear models (GLM) averaged according to the AAL template (25). To obtain these features, for each participant, we first fitted a GLM to the fMRI data during the MID task (see 10 for more details). Next, for each regressor in the GLM, the estimated coefficients were averaged according to the AAL template, producing one averaged coefficient for each ROI. ROIs for the left and right nucleus accumbens (NAcc) were further extracted according to a manually segmented MNI-152 brain (26) and added to the existing AAL ROIs, resulting in 118 ROIs. The following BOLD contrasts were included as features in our classification models to represent reward anticipation and consumption, respectively: 1) reward cue minus neutral cue and 2) reward outcome minus no-change outcome following reward cue. In addition, two striatal connectivity features emerging from (10) were included in our classification models, representing the psychophysiological interaction (PPI) under the reward outcome condition between 1) caudate and dorsal anterior cingulate cortex and 2) NAcc and mid-cingulate cortex. In total, 238 features (118 ROIs × 2 contrasts + 2 PPIs) were included in the classification models. All features were standardized to zero mean and unit variance before entered into the models.

### Classification and Importance-guided Sequential Model Selection

Two main classifiers were built to classify 1) MDD_Placebo_ vs. HC_Placebo_ and 2) MDD_Placebo_ vs. MDD_Amisulpride_. These were designed to capture features linked to 1) MDD, and 2) the effect of acute dopaminergic enhancement on whole-brain BOLD activation. To further test the hypothesis that dopaminergic enhancement transiently normalized reward-related abnormalities in MDD, a third classifier was built to classify MDD_Amisulpride_ vs. HC_Placebo_. Across analyses, we used logistic regression with elastic net regularization (27) as our classification models. The elastic net regularization has been shown to be well-suited for problems where the number of features is much greater than the number of observations (27). The models were trained and tested via the following nested cross-validation procedure. First, we performed model training on a development set containing 80% of the participants via a 3-fold grid search cross-validation procedure (stratified using class labels; Fig. 1b). Then, the model with the best regularization parameters was further tested on the evaluation set containing an independent set of 20% participants which the model had not seen during the training and validation phases. The above procedure was repeated 100 times to ensure stable performance was obtained on a large number of development-evaluation splits. The area under the receiver operating characteristics curve (AUC) was selected as the metric to quantify model performance.

To identify the set of most predictive features for each classifier (i.e., MDD_Placebo_ vs. HC_Placebo_ and MDD_Placebo_ vs. MDD_Amisulpride_), we adopted the following importance-guided sequential model selection procedure (Fig. 1a). Specifically, we first rank-ordered the features using the mean model weights across 100 implementations as a measure of predictability. Then, we built a series of truncated models such that each model only took the top k most predictive features as inputs to perform the classification tasks, with k varying from the top 1 most predictive feature to the number of participants involved in a given classifier. Imposing the number of participants as the upper limit was to ensure that models’ performance was not mainly driven by the regularization term. All truncated models underwent the nested cross-validation procedure described above and the test performance from each truncated model on the independent evaluation set was obtained. The set of features used by the truncated model achieving the highest AUC on the evaluation set were deemed as the optimal feature set.

### Statistical Analysis

The significance of the models’ performances against chance level was tested using a random permutation test scheme in which the truncated model based on the optimal feature set were re-trained on label shuffled training data (28). The entire test procedure was iterated 1000 times to empirically construct the null distribution of test AUCs. The p-values were obtained by comparing the AUC from the best truncated model based on unshuffled data against the empirical null distribution. The performances between models were statistically compared via Mann-Whitney U tests. Effect sizes between two distributions were calculated using Cohen’s d.

## Results

### Classification Performances

The best parsimonious models selected by the importance-guided model selection procedure (Fig. 1) based on most predictive features from whole-brain BOLD activations and striatal connectivity achieved high predictive performances (Table 2; see Fig. S1 for model performance as a function of top features). For both MDD_Placebo_ vs. HC_Placebo_ and MDD_Placebo_ vs.MDD_Amisulpride_, the AUC of the best parsimonious models were significantly above chance level (MDD_Placebo_ vs. HC_Placebo_: mean AUC = 0.87, *p* = 0.004; MDD_Placebo_ vs. MDD_Amisulpride_: mean AUC = 0.89, *p* = 0.002; Fig. 2a, b; Fig. S2). Compared with models trained using striatal features only (*Supplementary Methods*), the performances of the best parsimonious models based on whole-brain features were significantly better for both contrasts (*p*’s < 0.001, Mann-Whitney U test). The histograms of sum scores created by summing up the top feature values while taking into account the sign of the corresponding model weights demonstrated high separability between MDD_Placebo_ and HC_Placebo_ as well as between MDD_Placebo_ and MDD_Amisulpride_ (Fig. 2c, d). Overall, these results indicate that our models were able to extract highly predictive information embedded in the whole-brain BOLD signal.

**Table 2:** Classification performance for the best parsimonious models.

|  | MDD<sub>Placebo</sub> vs. |  | MDD<sub>Placebo</sub> vs. |  | MDD<sub>Placebo</sub> vs. |  | MDD<sub>Placebo</sub> vs. MDD<sub>Amisulpride</sub>: |  |
| --- | --- | --- | --- | --- | --- | --- | --- | --- |
|  | HC<sub>Placebo</sub> |  | MDD<sub>Amisulpride</sub> |  | HC<sub>Placebo</sub>: Striatum Only |  | Striatum Only |  |
|  | Mean | SD | Mean | SD | Mean | SD | Mean | SD |
| AUC | 0.87 | 0.12 | 0.89 | 0.09 | 0.59 | 0.14 | 0.61 | 0.17 |
| Accuracy | 0.77 | 0.12 | 0.80 | 0.10 | 0.59 | 0.13 | 0.59 | 0.13 |
| Sensitivity | 0.84 | 0.18 | 0.89 | 0.11 | 0.58 | 0.25 | 0.65 | 0.19 |
| Specificity | 0.72 | 0.22 | 0.67 | 0.24 | 0.59 | 0.22 | 0.50 | 0.28 |
| Number of Features | 48 |  | 44 |  | 6 |  | 11 |  |

**Figure 2:**
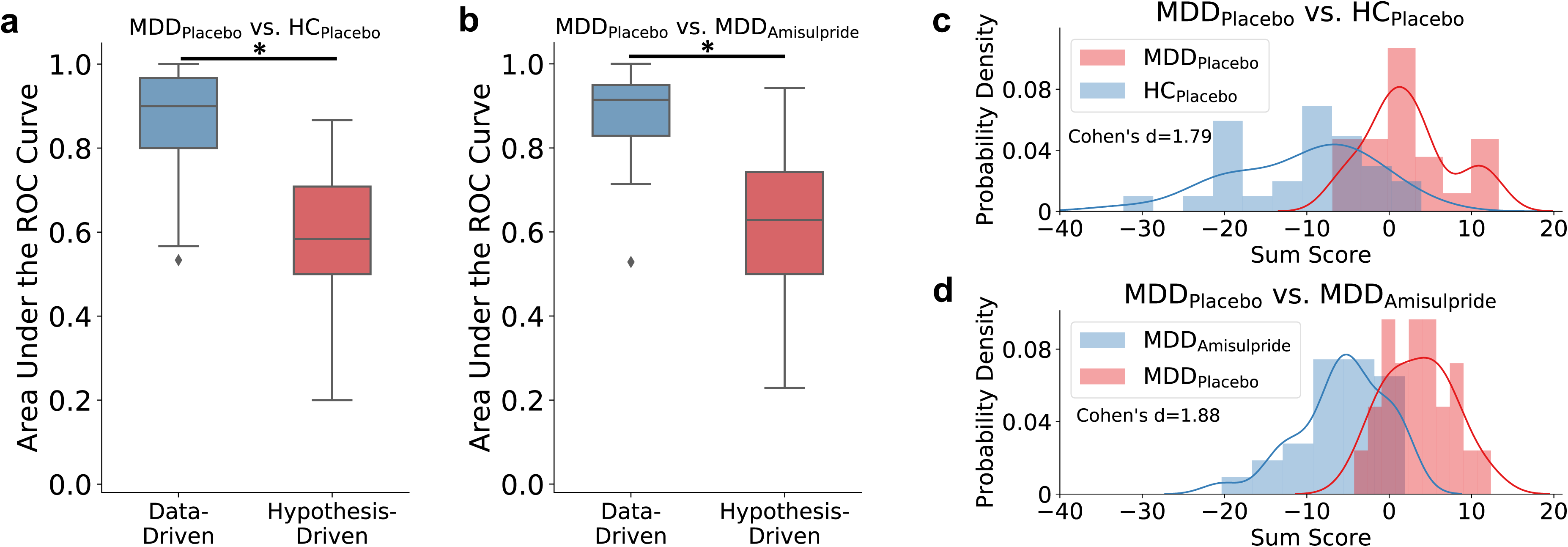
Comparing classification performance between the data-driven models based on features selected from the whole-brain and the hypothesis-driven models based only on striatal features for **a)** MDD_Placebo_ vs. HC_Placebo_ and **b)** MDD_Placebo_ vs. MDD_Amisulpride_ classifications. Asterisks denote significantly different median area under the Receiver Operating Characteristic (ROC) curve measures between the data-driven and hypothesis-driven models as assessed by the Mann-Whitney U test. The black markers denote outliers falling outside the ±1.5 interquartile range. The histogram of the signed sum score from the model-identified most predictive brain regions show high separability between **c)** MDD_Placebo_ vs. HC_Placebo_ and **d)** MDD_Placebo_ vs. MDD_Amisulpride_.

### Brain Regions Specific to Reward Anticipation

#### Positive model weights

The best parsimonious model for MDD_Placebo_ vs. MDD_Amisulpride_ identified the lateral orbitofrontal cortex (lOFC), visual cortex, anterior cingulate cortex (ACC), dorsomedial prefrontal cortex (dmPFC), mid-cingulate cortex (MCC), and precuneus as most predictive features with positive weights during reward anticipation (Fig. 3a; Table S1). This indicates that, within the MDD group, BOLD activation in these regions related to the contrast of reward cue minus neutral cue was reduced following administration of amisulpride compared to placebo. Critically, the lOFC, visual cortex, and MCC were also selected by the best MDD_Placebo_ vs. HC_Placebo_ model as top features having positive weights (Fig. 3b; Table S2), and at the same time these regions, except a right occipital region, were not among the most predictive features in the MDD_Amisulpride_ vs. HC_Placebo_ model (Fig. S3). Collectively, these findings indicate that, within the MDD group, amisulpride largely normalized the heightened BOLD activation in these regions toward reward cues. Other regions with positive weights in the MDD_Placebo_ vs. HC_Placebo_ classification included the thalamus, supplementary motor area (SMA), and the ventromedial prefrontal cortex (vmPFC). Again, these regions were not among the top features in the MDD_Amisulpride_ vs. HC_Placebo_ model (Fig. S3), suggesting that amisulpride mitigated the hyperactivation in these regions within the MDD group.

**Figure 3:**
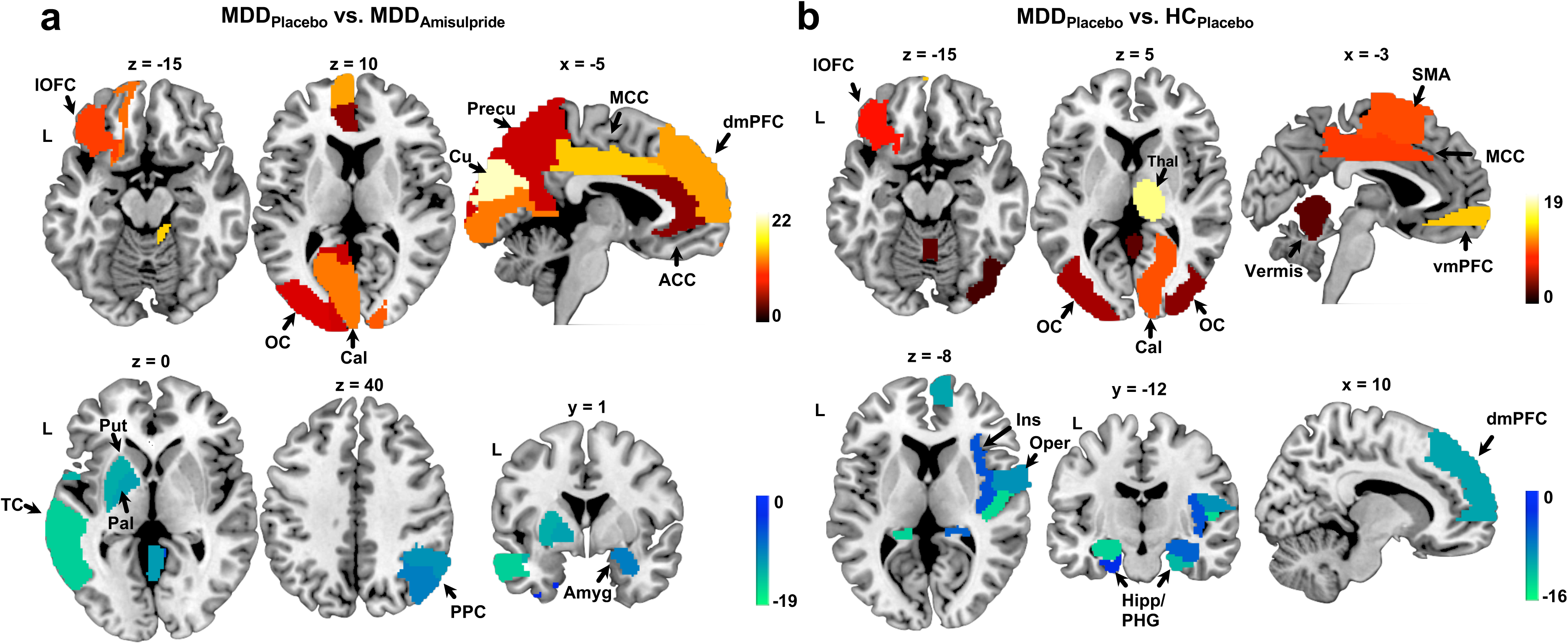
Weight maps showing the most predictive brain regions for the contrast of the reward minus neutral cue conditions. **a)** Weight map for the MDD_Placebo_ vs. MDD_Amisulpride_ model. Positive weights indicate higher BOLD in the MDD_Placebo_ group relative to the MDD_Amisulpride_ group and negative weights indicate the opposite direction. **b)** Weight map for the MDD_Placebo_ vs. HC_Placebo_ model, with positive weights indicating higher BOLD in the MDD_Placebo_ group relative to the HC_Placebo_ group and vice versa. ACC: anterior cingulate cortex; Amyg: amygdala; Cal: calcarine sulcus; Cu: cuneus; dmPFC: dorsomedial prefrontal cortex; Hipp: hippocampus; Ins: insula; lOFC: lateral orbitofrontal cortex; MCC: middle cingulate cortex; OC: occipital cortex; Oper: operculum; Pal: pallidum; PHG: parahippocampal gyrus; PPC: posterior parietal cortex; Precu: precuneus; Put: putamen; SMA: supplementary motor area; TC: temporal cortex; vmPFC: ventromedial prefrontal cortex.

#### Negative model weights

Regions selected by the best MDD_Placebo_ vs. MDD_Amisulpride_ model with negative model weights included the putamen, pallidum, amygdala, posterior parietal cortex (PPC), and temporal cortex (Fig. 3a; Table S1). The negative weights observed in the putamen and pallidum were consistent with the hypothesis that amisulpride might have increased dopaminergic signaling in the basal ganglia in MDD (10, 14). This effect is rather pronounced as the MDD_Amisulpride_ vs. HC_Placebo_ model showed that the contrast of reward cue minus neutral cue evoked higher activation in the putamen in the MDD_Amisulpride_ group even compared with the HC_Placebo_ group (Fig. S3). Within the MDD_Placebo_ group, reduced activation in the operculum, hippocampus, parahippocampal gyrus (PHG), and dmPFC was observed relative to HCs during reward anticipation (features in the MDD_Placebo_ vs. HC_Placebo_ model with negative weights; Fig. 3b; Table S2). The reduced activation in the hippocampus and operculum persisted in the MDD_Amisulpride_ vs. HC_Placebo_ model (Fig. S3), indicating that amisulpride had limited effects in these regions.

### Brain Regions Specific to Reward Consumption

#### Positive model weights

Examining features selected from the contrast of reward minus no change outcomes in the MDD_Placebo_ vs. MDD_Amisulpride_ model revealed that the lOFC, PPC, superior frontal gyrus, and the pre- and post-central gyrus were selected as most predictive features with positive weights (Fig. 4a, Table S3). This indicates reduced activation in these regions during reward consumption in MDD_Amisulpride_ compared with MDD_Placebo_. Of note, the lOFC and PPC emerged as among the most predictive features with positive weights in the MDD_Placebo_ vs. HC_Placebo_ model (Fig. 4c, Table S4). Additionally, while the lOFC hyperactivation was still observed in the MDD_Amisulpride_ vs. HC_Placebo_ model, the PPC was not identified as a predictive feature (Fig. S3). Overall, these results suggest that, under placebo, the MDD group was characterized by increased BOLD activity in these regions during reward consumption relative to HC and that the hyperactivation was reduced by amisulpride. Other brain regions identified as most predictive features with positive weights in the MDD_Placebo_ vs. HC_Placebo_ model included the inferior frontal gyrus, PCC, precuneus, and MCC. The lack of predictability from these regions between MDD_Amisulpride_ and HC_Placebo_ (Fig. S3) again suggests a mitigating effect of amisulpride on the hyperactivation in these regions.

**Figure 4:**
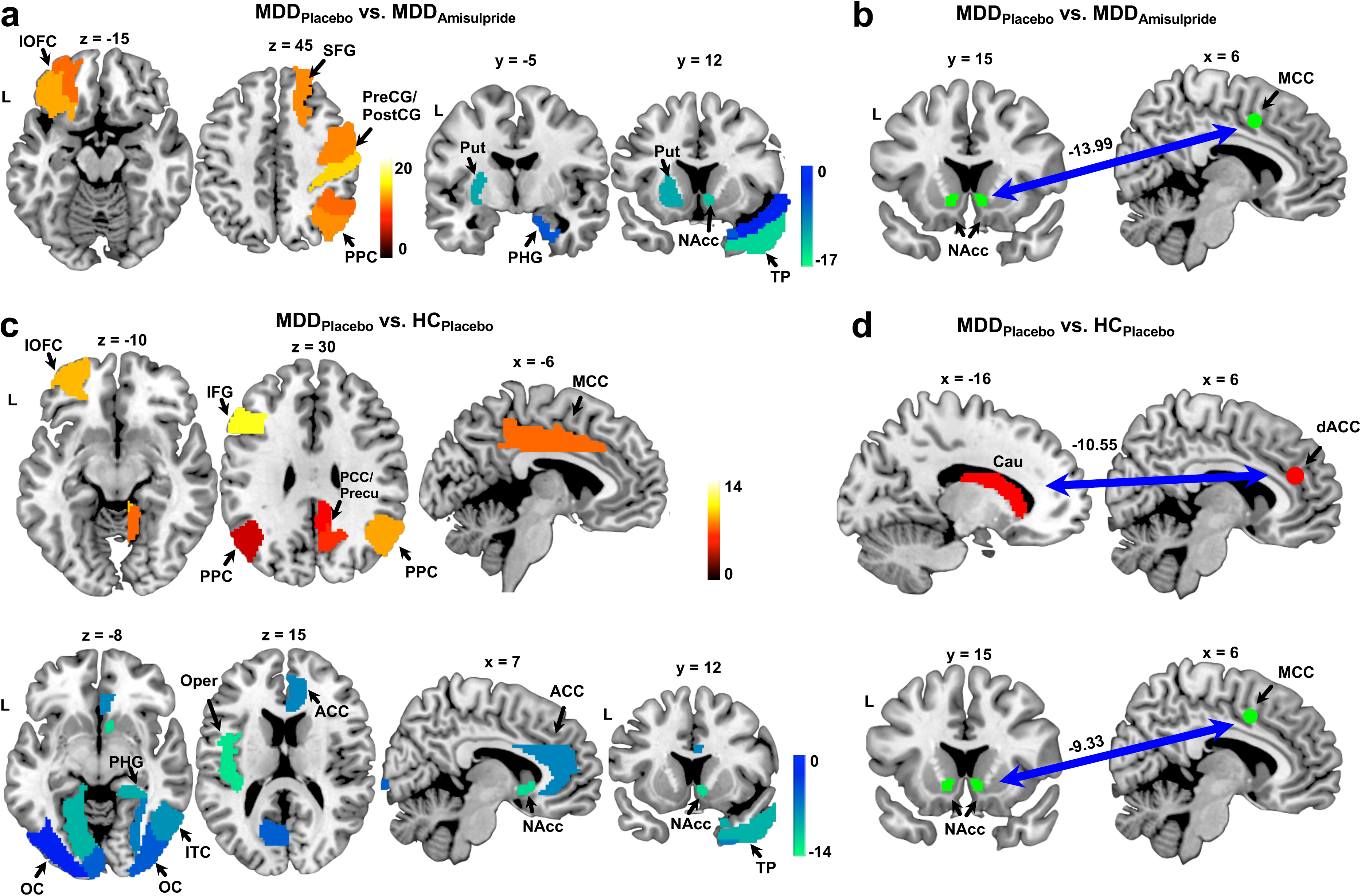
Weight maps showing the most predictive brain regions/connectivity for the contrast of reward minus no-change outcomes. **a)** Weight map for the MDD_Placebo_ vs. MDD_Amisulpride_ model, with positive weights indicating higher BOLD in the MDD_Placebo_ group relative to the MDD_Amisulpride_ group and vice versa. **b)** Negative weight assigned to the NAcc-MCC connectivity in the MDD_Placebo_ vs. MDD_Amisulpride_ model. **c)** Weight map for the MDD_Placebo_ vs. HC_Placebo_ model. Positive weights indicate higher BOLD in the MDD_Placebo_ group relative to the MDD_Amisulpride_ group and vice versa. **d)** Negative weights assigned to the Caudate-dACC and NAcc-MCC connectivity features by the MDD_Placebo_ vs. HC_Placebo_ model. Abbreviations followed those used in Fig. 2. Cau: caudate; dACC: dorsal anterior cingulate cortex; IFG: inferior frontal gyrus; ITC: inferior temporal cortex; PCC: posterior cingulate cortex; PreCG/PostCG: pre- and post-central gyrus; SFG: superior frontal gyrus; TP: temporal pole.

#### Negative model weights

The most predictive regions from the contrast of reward minus no change outcomes with negative weights in the MDD_Placebo_ vs. MDD_Amisulpride_ model included the putamen, NAcc, PHG, and temporal pole (Fig. 4a, Table S3), as well as the connectivity between the NAcc and MCC (Fig. 4b). This suggests that, within the MDD group, amisulpride increased BOLD activation and corticostriatal connectivity to reward feedback in these regions. Highlighting again convergence, the NAcc, PHG, temporal pole, and the NAcc-MCC connectivity were also selected as most predictive features having negative weights in the MDD_Placebo_ vs. HC_Placebo_ classification (Fig. 4c, d, Table S4), and none of these regions was selected as among the top predictive features in the MDD_Amisulpride_ vs. HC_Placebo_ model (Fig. S3). Thus, in MDD, amisulpride normalized both hypoactivation and hypoconnectivity in response to rewards in these regions. Other most predictive features with negative weights in the MDD_Placebo_ vs. HC_Placebo_ model included the visual cortex, inferior temporal cortex, operculum, ACC, and the connectivity between the caudate and dACC. These features, except the caudate-dACC connectivity, were not identified as among the top features in the MDD_Amisulpride_ vs. HC_Placebo_ model (Fig. S3), indicating increased activation to rewards in these regions following amisulpride administration in the MDD group. The fact that amisulpride did not normalize the hypoconnectivity between caudate and dACC in the MDD group is consistent with previously published ROI-based results obtained on a subset of the participants (10).

## Discussion

This study utilized a machine learning based approach (importance guided sequential variable selection using logistic regression with elastic net regularization; Fig. 1) to identify reliable features across the entire brain that distinguished unmedicated individuals with MDD from HC (under placebo condition) as well as features linked to normalization of putative MDD-related abnormalities after an acute pharmacological challenge hypothesized to increase dopaminergic signaling along mesolimbic pathways. In addition to increased striatal activation in the MDD_Amisulpride_ relative to MDD_Placebo_ group (which is consistent with ROI-based conventional analyses of a smaller subset of the participants included here, 10), the classification model also identified an extensive set of brain regions differentiating individuals with MDD receiving amisulpride vs. placebo. Notably, the classification performance from models relying on whole-brain features was significantly higher than the performance of models based only on striatal features, indicating that a pharmacological challenge that putatively increased dopaminergic signaling induced widespread differential BOLD activity among depressed individuals during reward processing. Comparing the most predictive brain regions between the MDD_Placebo_ vs. MDD_Amisulpride_ model and the MDD_Placebo_ vs. HC_Placebo_ model suggested that amisulpride had a bidirectionally normalizing effect on reward-related activation and functional connectivity of brain regions spanning the lOFC, NAcc, PHG, MCC, PPC, and areas of the visual cortex among depressed individuals. Taken together, these results highlight the unique contribution of machine learning-based approaches to examine pharmacologically-induced effects across the entire brain when assessing circuit engagement by targeted treatment options for mood and psychiatric disorders. The study provides novel evidence for the mechanism through which (transient) dopaminergic enhancement might restore system-level activity during reward processing among individuals with MDD.

Amisulpride appeared to have bi-directional normalizing effects on brain activation and functional coordination among depressed individuals. Within the striatum, consistent with previous ROI-based analyses based on a subset of the participants used here (10), results from our classification models showed that decreased striatal/basal ganglia activation and corticostriatal connectivity among depressed individuals were enhanced following acute administration of amisulpride. This supports the validity of the importance-guided model selection procedure and fits the view that lower doses of amisulpride enhance dopaminergic signaling in the striatum (17).

Among regions outside the striatum, one notable finding was that increased lOFC activation during reward anticipation in MDD was reduced after administration of amisulpride. Neurophysiological evidence has shown that subpopulations of neurons in the lOFC respond to non-reward/unpleasant events and maintain elevated firing rate after such events (29). This led to the theory implicating overly reactive and prolonged activation of the lOFC non-reward circuit as a potential mechanism underlying depression (30). Previous studies have documented increased lOFC activation in MDD (31), and our result fits this theoretical view. In the MDD_Amisulpride_ group, reduced lOFC activation suggests that amisulpride may normalize reward processing by decreasing lOFC hyperactivation, consistent with previous reports that improvements in depressive symptoms were accompanied by reduced lOFC activation (32) and electrical stimulation of the lOFC acutely improved depressive symptoms (33).

In addition to effects in frontostriatal circuitry, amisulpride restored hypoactivation in the parahippocampal gyrus and temporal pole in MDD. The hippocampus and parahippocampal complex connect with the medial OFC and are hypothesized to facilitate the formation of episodic memory regarding reward (34). Decreased hippocampal activation has emerged in MDD and prolonged/repeated depressive episodes have been linked to reduced hippocampal volume (35, 36). These abnormalities have been linked to dysfunctions in both memory encoding and retrieval characteristic of MDD, even after treatment (37, 38). The fact that amisulpride restored parahippocampal and temporal pole activation suggests that interventions aiming to increase dopaminergic signaling might improve encoding and retrieval of positive memories in MDD. However, it should be noted that hippocampal activation did not differentiate between the MDD_Amisulpride_ and MDD_Placebo_ group, suggesting that the effects on memory might be limited following a single acute pharmacological challenge.

Hyperactivation in the mid-cingulate cortex towards the reward cue was also reduced among depressed individuals after amisulpride. Moreover, amisulpride also reduced reward cue-evoked activations in adjacent ACC and dmPFC. The supracallosal part of the cingulate cortex receives neuronal projections from the lOFC and is thought to also encode non-reward and punishing events such as physical and social pain (39, 40). A recent study has identified a nociceptive pathway between the mid-cingulate cortex and posterior insula responsible for generating a hypersensitive state for pain, providing a mechanism for the increased pain sensitivity by psychosocial factors (41). The reduced hyperactivation in these regions following amisulpride administration may indicate decreased sensitivity to negative affective states among individuals with MDD and therefore priming or biasing them toward reward.

In MDD, amygdalar activation evoked by reward cues was enhanced following amisulpride. Reduced amygdalar response to positive and rewarding stimulus, coupled with heightened amygdalar activation toward negative stimulus, are well-documented findings in MDD, which highlights an imbalanced reactivity toward emotionally-salient cues (42). Antidepressant treatment has been shown to address this imbalance by partially normalizing the bi-directional abnormal amygdalar activation (42, 43). These findings were further bolstered by the recent report that enhanced amygdalar response toward positive memories through real-time fMRI neurofeedback was associated with reduction in depressive symptoms (44). The increased amygdalar activation evoked by reward cues is consistent with these studies and implicates improved sensitivity toward reward following acute dopaminergic enhancement.

It should be noted that while several regions showed predictive power following the administration of amisulpride, it is difficult to assess whether changes in these regions reflected a direct modulation resulting from the enhanced dopaminergic signaling or alternatively reflected secondary responses through network interactions. Future studies could utilize network analysis and/or neural perturbation methods to further dissociate direct vs. indirect effects (33). In addition, amisulpride also has 5-HT_7_ antagonism (45), which has been hypothesized to contribute to its antidepressant property. While we cannot rule out that the effects observed here may be partially caused by this off-target mechanism, additional research is needed to distinguish the effect of dopaminergic enhancement vs. 5-HT_7_ antagonism of amisulpride. Lastly, we only focused on investigating the effects of dopaminergic enhancement on reward processing among depressed individuals. Future studies could seek to examine the effect of enhanced dopamine on whole-brain fMRI activity in depression under additional conditions.

## Acknowledgements

This project was supported by R01 MH068376, R37 MH068376 and R01MH095809 from the National Institute of Mental Health (Dr. Pizzagalli). The content is solely the responsibility of the authors and does not necessarily represent the official views of the National Institutes of Health.

## Author Contributions

DAP designed both studies study and obtained funding for both; RC, MB, FG and GV collected data; YL, RA, ELB, RHK performed the analyses; YL, DAP, RA, and PA wrote the manuscript. All authors approved the manuscript.

## Data Availability

Data are available at the NIMH Data Archive (https://nda.nih.gov/)

## Code Availability

Analysis scripts are available upon request.

