## Supplemental Materials for "Machine learning identifies large-scale reward-related activity modulated by dopaminergic enhancement in major depression"

### Supplementary Material

### Supplementary Methods

#### Participants

Across the two studies, additional exclusion criteria included: serious or unstable medical illness (e.g., cardiovascular, endocrine, neurologic or hematologic disease); renal insufficiency, clinical or laboratory evidence of hypothyroidism; history of seizure disorder, cocaine or stimulant use (e.g., amphetamine, cocaine, methamphetamine), dopaminergic drug use (including methylphenidate), dementia, or adverse drug reactions or allergy to amisulpride; pregnancy; use of oral contraceptives or hormone therapy in the previous six months; MRI safety contraindications; severe concussion or loss of consciousness (>2 min). For the MDD group, additional exclusion criteria included: suicidal ideation; any psychotropic medication in the prior two weeks (six weeks for fluoxetine; six months for dopaminergic drugs or neuroleptics); prior electroconvulsive therapy; and a history or current diagnosis of any of the following DSM-IV psychiatric illnesses: organic mental disorder, schizophrenia, schizoaffective disorder, delusional disorder, psychotic disorders not otherwise specified, bipolar disorder, mood congruent or mood incongruent psychotic features, lifetime substance dependence, substance abuse within the last 12 months (except cocaine or stimulant abuse, which led to exclusion). Selected anxiety disorders (simple phobia, social anxiety disorder and generalized anxiety disorders) were allowed only if secondary to MDD.

Participants deemed to be eligible after a phone screen were invited for a clinical evaluation that included an interview to further assess medical and neurological history; administration of the Structured Clinical Interview for DSM-IV-TR Axis I Disorders, Research Version, Non-patient Edition (SCID-I/NP)^1^; and several self-report scales, including the Beck Depression Inventory (BDI-II)^2^, the Mood and Anxiety Symptom Questionnaire (MASQ)^3^, and Snaith–Hamilton Pleasure Scale (SHAPS)^4^. For the imaging session, in order to avoid craving effects, participants were asked to consume their usual amount of caffeine and/or nicotine. The MDD and HC groups did not differ in their smoking status but showed a significant difference in their caffeine consumption (p = 0.002; Table S5).

#### Rationale for Using Amisulpride

Amisulpride – a dopamine D2 receptor antagonist – was selected due to its high affinity for D_2_/D_3_ dopamine receptors, particularly in the mesocorticolimbic dopaminergic pathway^5^. At low doses, amisulpride has been found in preclinical studies to preferentially blocks presynaptic dopamine autoreceptors, leading to increased striatal dopamine release and prohedonic effects^6–10^. In humans, the 50 mg dose used in the current study has been found to have no effects on mood or sensory-motor coordination when given as a single dose^11^, whereas it was found to have antidepressant and anti-anhedonic effects in depressive disorders when given over weeks^12–14^. Note that, although prior preclinical and clinical findings using low doses of amisulpride are consistent with presynaptic autoreceptor blockade, alternative interpretations exist, including (1) feedback excitation of dopamine neurons, likely via blockade of somatodendritic autoreceptors^15^ and (2) postsynaptic D2 blockade leading to feedback excitation of DA neurons^16^.

#### The Monetary Incentive Delay Task

At the beginning of each trial a visual cue was presented for 0.5 sec to indicate the trial type (reward: +$; penalty: –$; no incentive: 0$). After a jittered inter-stimulus interval (2.25–3.75 sec), a red target square was presented for 0.15 sec, and participants were instructed to press a button as fast as possible after seeing the square. After a jittered delay (2.4–3.9 sec), a visual feedback was presented for 1.25 sec to inform the participant about the outcome (reward, penalty, or no change). Trials were separated by a third jittered interval (1.5–4.5 sec). Participants were instructed that fast responses to the red square would increase their chances to receive rewards and avoid penalties. Task difficulty was calibrated by using the 70th percentile of each participant’s reaction time during a practice session as the threshold for success. In reward trials, success led to monetary gains ($1.96 to $2.34), whereas slow responses led to no-change. In penalty trials, success led to no-change, whereas slow responses led to monetary penalties (-$1.81 to -$2.19). No-incentive trials always ended with no-change feedback. For both studies, five runs of 24 trials (8 for each trial types) were administered, and no feedback about cumulative earnings was provided.

#### Data Acquisition

For both studies, MRI data were acquired at the McLean Imaging Center using a Siemens Tim Trio 3T MR scanner equipped with a 32-channel head coil. Functional data (461 volumes) were collected using a T_2_-weighted spin echo planar imaging sequence (TR: 3000ms; TE: 30ms; FOV: 224mm; matrix: 64x64; resolution: 3.5x3.5x2.0mm; number of slices: 57 aligned to the AC–PC plane). For co-registration purpose, high-resolution T_1_-weighted MPRAGE images were acquired (TR: 2200ms; TE: 1.54ms; FOV: 230mm; matrix: 192x192; resolution: 1.22mm^3^; number of slices: 144].

#### MRI Preprocessing

fMRI preprocessing is described in detail elsewhere^17^. In brief, fMRI data preprocessing relied on Statistical Parametric Mapping (SPM12; Wellcome Department of Cognitive Neurology, London, UK), and included coregistration of functional and anatomical images, segmentation, nonlinear volume-based spatial normalization (Montreal Neurological Institute (MNI) template), and spatial smoothing with a Gaussian filter (6mm FWHM). Artifact Detection Tools (ART) (http://web.mit.edu/swg/software.htm) was used to identify and exclude outlier time points in the global mean image time series (threshold: 3 standard deviations (SD) from the mean) and movement (threshold: 0.7mm; measured as scan-to-scan movement, separately for translation and rotation) parameters.

#### Classification Based on Striatal Features

Prior literature demonstrated that MDD patients show reduced activation in the striatum during the MID task^18^ and such striatal blunting could be rescued by amisulpride^17^. To obtain model performances based on these known striatal features as a comparison, we repeated the above 2 classification tasks via the sequential model selection procedure using only the striatal ROI (bilateral caudate, putamen, and NAcc) and connectivity (2 striatal psychophysiological interaction) features^17^. The performances from these hypothesis-driven models were used as a benchmark for the whole-brain classification model.

### Supplementary Results

#### Figure S1

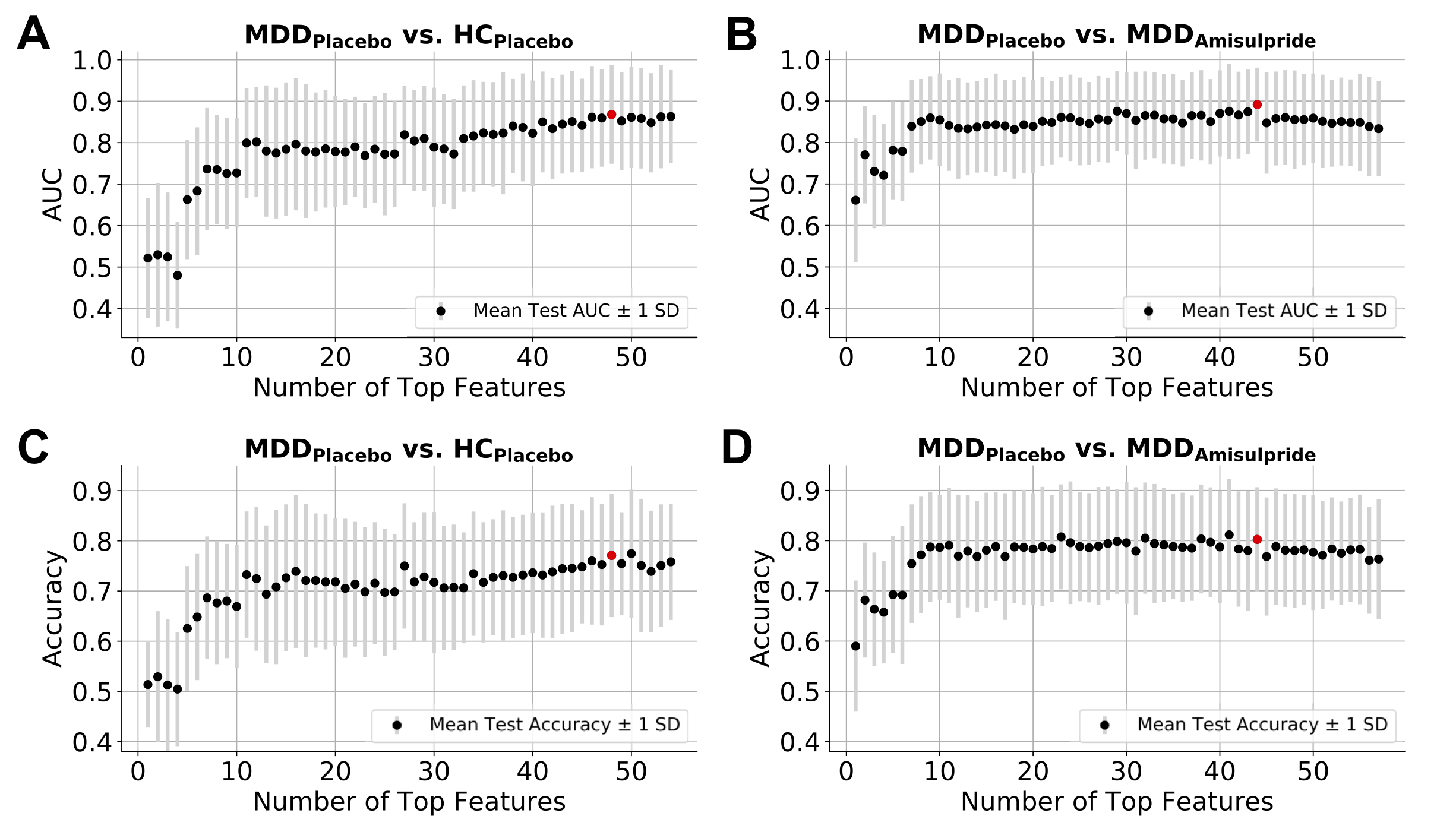

**Figure S1**: Classification performance as a function of number of top features. The AUCs for the truncated models are shown for **A)** MDD_Placebo_ vs. HC_Placebo_ classification and **B)** MDD_Placebo_ vs. MDD_Amisulpride_ classification. The red dot denotes the highest AUC achieved. The models’ accuracy scores are shown in **C)** and **D)**, respectively, with the red dot denoting the corresponding accuracy score at the number of features achieving the highest AUC.

#### Figure S2

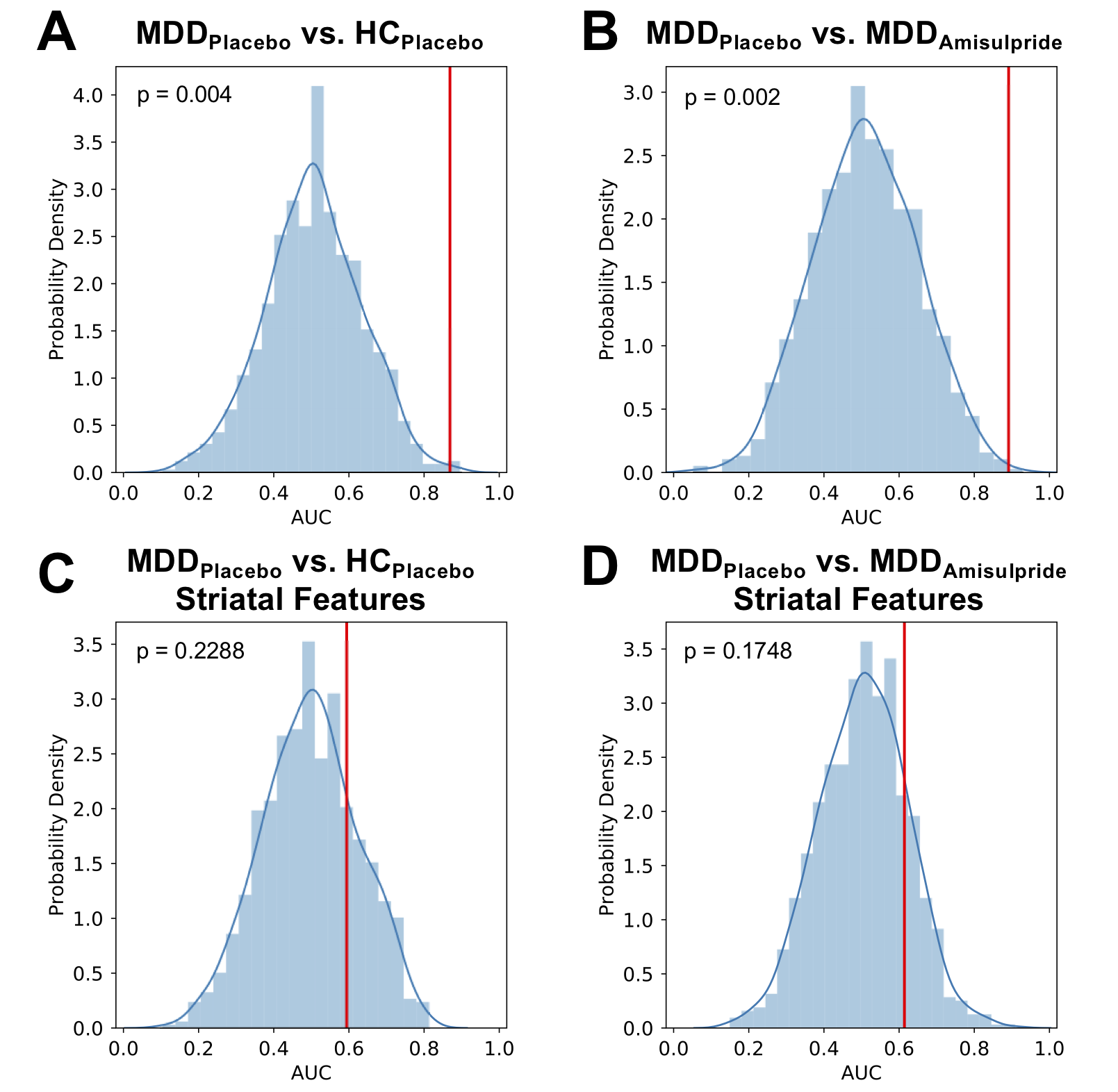

**Figure S2**: Permutation test results for the best parsimonious model in **A)** MDD_Placebo_ vs. HC_Placebo_ classification and **B)** MDD_Placebo_ vs. MDD_Amisulpride_ classification. The test results for the corresponding classification models based on striatal features only are shown in **C)** and **D)**, respectively. The histogram represents the empirical null distribution of AUCs from 1000 permutation iterations. The red vertical line marks the AUC obtained from the actual unshuffled data.

#### Figure S3

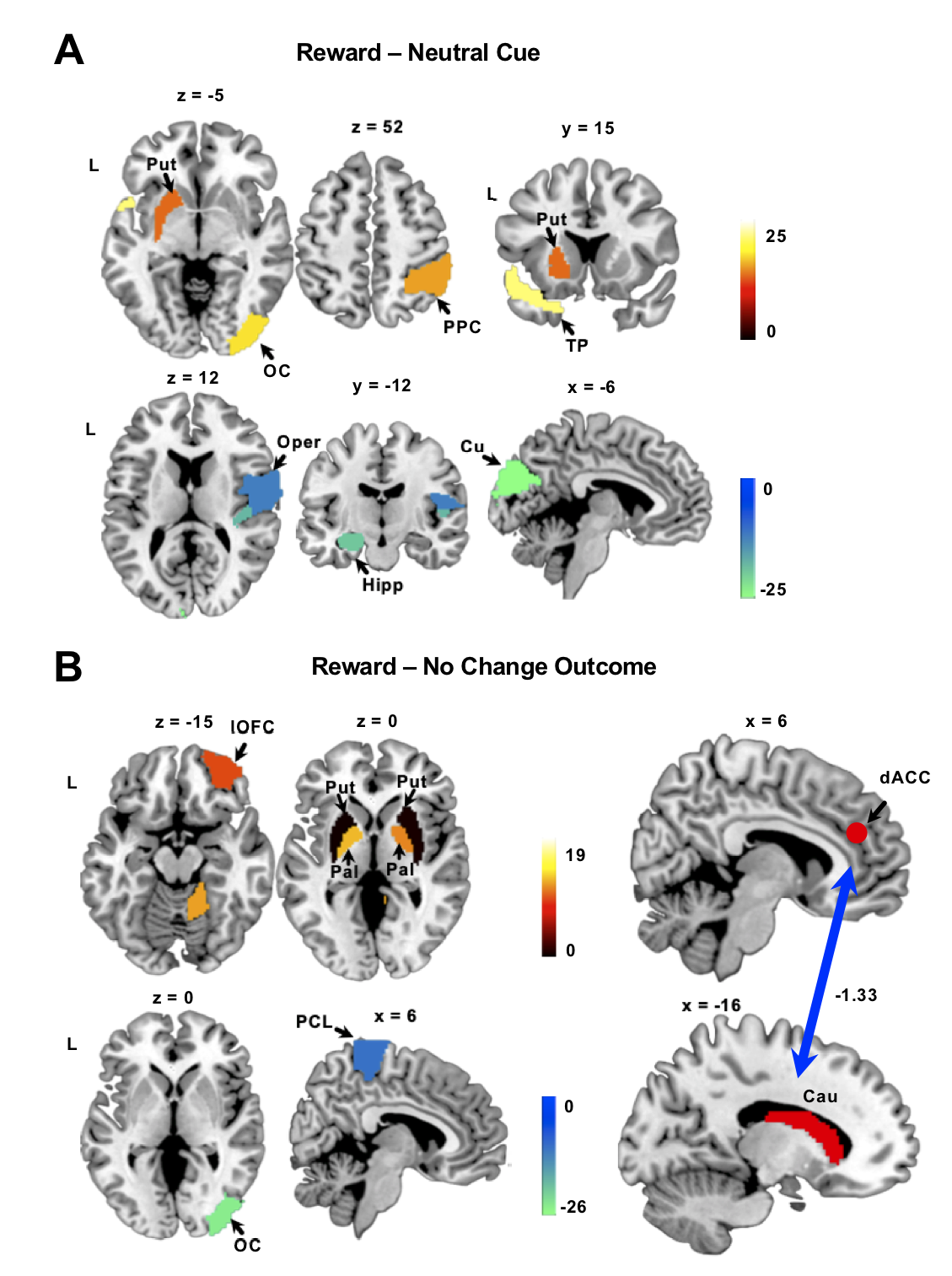

**Figure S3**: Weight map showing the most predictive brain regions and connectivity in the MDD_Amisulpride_ vs. HC_Placebo_ model from **A)** the reward minus neutral cue condition and **B)** the reward minus no-change outcome condition. Positive weights indicate higher BOLD/connectivity in the MDD_Amisulpride_ group relative to the HC_Placebo_ group and vice versa. Abbreviations follow those used in Figs. 2 and 3. PCL: paracentral lobule.

#### Table S1

MDD_Placebo_ vs. MDD_Amisulpride_: Top features from the Reward – Neutral Cue condition

| **Top Feature** | **AAL ROI Label** | **Model Weight** |
| --- | --- | --- |
| Occipital/Visual Cortex | Cuneus_L | -20.65 |
|  | Occipital_Inf_R | 17.25 |
|  | Calcarine_L | -11.94 |
|  | Cuneus_R | -10.85 |
|  | Occipital_Mid_L | -6.7 |
|  | Occipital_Sup_L | -6.36 |
| Middle temporal Cortex | Temporal_Mid_L | 15.96 |
| Mid-cingulate Cortex | Cingulum_Mid_L | -14.68 |
| Temporal Pole | Temporal_Pole_Sup_L | 13.64 |
|  | Temporal_Pole_Mid_L | 2.84 |
| Dorso-medial PFC | Frontal_Sup_Medial_L | -13.29 |
| Putamen | Putamen_L | 13.28 |
| Pallidum | Pallidum_L | 12.28 |
| Lateral OFC | Frontal_Sup_Orb_L | -11.51 |
|  | Frontal_Inf_Orb_L | -9.93 |
| Posterior Parietal Cortex | Parietal_Inf_R | 11.28 |
|  | Angular_R | 9.72 |
| Amygdala | Amygdala_R | 9.65 |
| Precuneus | Precuneus_L | -6.16 |
| Anterior Cingulate Cortex | Cingulum_Ant_L | -4.18 |
| Cerebellum | Vermis_8 | 18.84 |
|  | Cerebellum_3_R | -14.9 |
|  | Cerebellum_7B_R | -13.18 |
|  | Vermis_6 | 12.16 |
|  | Vermis_4_5 | 11.92 |
|  | Cerebellum_Crus1_R | -7.77 |
|  | Cerebellum_4_5_R | 5.66 |

#### Table S2

MDD_Placebo_ vs. HC_Placebo_: Top features from the Reward – Neutral Cue condition

| **Top Feature** | **AAL ROI Label** | **Model Weight** |
| --- | --- | --- |
| Thalamus | Thalamus R | 18.69 |
| Ventral Medial PFC | Frontal_Med_Orb_L | 14.17 |
| Insula/Operculum | Heschl_R | -13.35 |
|  | Rolandic_Oper_R | -8.15 |
|  | Insula_R | -3.62 |
| Hippocampus | Hippocampus_L | -12.85 |
|  | Hippocampus_R | -4.73 |
| Parahippocampal Gyrus | Parahippocampal_R | -10.92 |
|  | Parahippocampal_L | -0.98 |
| Occipital/Visual Cortex | Calcarine_R | 10.43 |
|  | Occipital_Mid_L | 5.02 |
|  | Occipital_Mid_R | 4.2 |
|  | Occipital_Inf_R | 2.03 |
| Supplementary Motor Area | Supp_Motor_Area_L | 10.26 |
| Mid-cingulate Cortex | Cingulum_Mid_L | 9.69 |
| Dorso-medial PFC | Frontal_Sup_Medial_R | -9.51 |
| Lateral OFC | Frontal_Inf_Orb_L | 8.37 |
| Cerebellum | Vermis_8 | -12.3 |
|  | Vermis_4_5 | 2.77 |

#### Table S3

MDD_Placebo_ vs. MDD_Amisulpride_: Top features from the Reward – No Change Outcome condition

| **Top Feature** | **AAL ROI Label** | **Model Weight** |
| --- | --- | --- |
| Temporal Pole | Temporal_Pole_Mid_R | -15.65 |
|  | Temporal_Pole_Sup_R | -2.48 |
| Nucleus Accumbens | NAcc_R | -14.31 |
| Accumbens - MCC Connectivity |  | -13.99 |
| Postcentral Gyrus | Postcentral_R | 12.89 |
| Putamen | Putamen_L | -12.47 |
| Lateral OFC | Frontal_Inf_Orb_L | 11.45 |
|  | Frontal_Mid_Orb_L | 9.37 |
| Posterior Parietal Cortex | Parietal_Sup_R | 10.62 |
|  | Angular_R | 10.48 |
|  | Parietal_Inf_R | 9.24 |
| Superior Frontal Gyrus | Frontal_Sup_R | 10.42 |
| Precentral Gyrus | Precentral_R | 10.22 |
| Parahippocampal Gyrus | Parahippocampal_R | -6.28 |
| Cerebellum | Cerebellum_9_R | 12.5 |
|  | Cerebellum_7B_R | 12.25 |
|  | Vermis_8 | -4.01 |

#### Table S4

MDD_Placebo_ vs. HC_Placebo_: Top features from the Reward – No Change Outcome condition

| **Top Feature** | **AAL ROI Label** | **Model Weight** |
| --- | --- | --- |
| Pallidum | Pallidum_L | 12.67 |
| Insula/Operculum | Rolandic_Oper_L | -12.57 |
| Nucleus Accumbens | NAcc_R | -12.01 |
| Inferior Frontal Gyrus | Frontal_Inf_Tri_L | 11.11 |
| Caudate - dACC Connectivity |  | -10.55 |
| Occipital/Visual Cortex | Lingual_L | -9.81 |
|  | Fusiform_R | -6.71 |
|  | Calcarine_L | -4.96 |
|  | Occipital_Inf_R | -4.79 |
|  | Occipital_Inf_L | -1.39 |
| Temporal Pole | Temporal_Pole_Mid_R | -9.64 |
| Accumbens - MCC Connectivity |  | -9.33 |
| Parahippocampal Gyrus | Parahippocampal_R | -9.27 |
| Lateral OFC | Frontal_Mid_Orb_L | 9.26 |
| Posterior Parietal Cortex | Angular_R | 8.81 |
|  | Parietal_Sup_R | 7.9 |
|  | Parietal_Inf_R | 6.94 |
|  | Angular_L | 4.24 |
| Inferior Temporal Cortex | Temporal_Inf_R | -7.84 |
| Mid-cingulate Cortex | Cingulum_Mid_L | 7.52 |
| Anterior Cingulate Cortex | Cingulum_Ant_R | -7.08 |
| Precuneus/Posterior Cingulate Cortex | Precuneus_R | 6.25 |
|  | Cingulum_Post_R | 5.2 |
| Cerebellum | Cerebellum_8_L | -10.06 |
|  | Cerebellum_9_R | 9.91 |
|  | Cerebelum_Crus2_R | 9.69 |
|  | Cerebellum_3_R | 9.57 |
|  | Cerebellum_7b_R | 8.11 |
|  | Cerebellum_4_5_R | 7.48 |

#### Table S5

Smoking status and caffeine consumption

|  | MDD_Amisulpride_ | | MDD_Placebo_ | | HC_Placebo_ | |
| --- | --- | --- | --- | --- | --- | --- |
|  | N | % | N | % | N | % |
| Current smokers^a^ | 1 | 3.2 | 0 | 0 | 0 | 0 |
| Past smokers^a^ | 1 | 3.2 | 3 | 11.5 | 0 | 0 |
|  | Mean | SD | Mean | SD | Mean | SD |
| Caffeine consumption (mg/day)^b^ | 92.6 | 72.7 | 143.9 | 95.2 | 66.8 | 51.2 |

^a^ No difference in smoking status was observed across the groups (p’s > 0.13).

^b^ A significant difference in daily caffeine consumption across the groups was observed via a one-way ANOVA (p = 0.002).
